# *In situ* near-infrared non-destructive monitoring of sugar accumulation reveals that single berries ripening takes only 20 days

**DOI:** 10.1101/2024.12.10.627588

**Authors:** Flora Tavernier, Elias Motelica-Heino, Miguel Thomas, Theresa Herbold, Mengyao Shi, Loïc Le Cunff, Charles Romieu, Vincent Segura

## Abstract

Understanding how climate change impacts berry ripening physiology is essential for selecting genotypes that balance sugars and acids under warming conditions. In this context, we used a portable near-infrared spectrometer in the vineyard, to monitor sugar and acid evolution in individual berries from 10 grapevine varieties over two years. Spectra were periodically acquired on the same berries all along ripening, and a subset of these berries was also collected for sugars and organic acids quantification by HPLC, to train partial least square regression models. Prediction models for glucose, fructose, and malic acid concentrations were characterized by validation R² of 0.71, 0.64, and 0.55, respectively. We further used these models to follow sugar accumulation in individual berries and observed that single berries ripen two times faster than found in samples composed of multiple berries. Our results pave avenues toward precise quantitative approaches on sugar and acid fluxes in berry ripening studies.

## Introduction

Climate change poses an increasing threat to plants in general and, in particular, grapevine (IPCC, 2023). Increasing temperatures advances its phenological development, leading to ripening under hotter and drier conditions. These changes affect not only growth but also berry composition, leading to higher sugar concentration and therefore potential alcohol, while reducing acidity, which are critical enological variables for wine quality (Castellarin et al., 2007; Gambetta et al., 2020; Bécart et al., 2022).

Grape development follows a double sigmoid curve, segmented into two main phases: the green stage and ripening (Coombe and Hale, 1973; Coombe and McCarthy, 2000). The green stage corresponds to cell division and expansion, related to the accumulation of malic and tartaric acids as major osmotica (Ojeda et al., 1999; Terrier and Romieu, 2001). The transition between the green stage and ripening is marked by veraison, when the berries rapidly soften and start to accumulate sugars at high-rate, before anthocyanin synthesis begins and provides color to pigmented-skin varieties (Lund and Bohlmann, 2006). Berry growth resumes briefly after softening, doubling their volume through water absorption until sugar accumulation stops and the berries begin to shrivel (Coombe and Bishop, 1980; Savoi et al., 2021; Daviet et al., 2023). Temperature, rainfall, and photosynthetically active radiation (PAR), differentially impact the accumulation of water and sugar in berries thus influencing solute concentration (Suter et al., 2021). The same authors also reported that the duration of the sugar accumulation period ranges between 1 and 2 months according to the genotype. This duration is often underestimated because measurements typically begin at mid-coloring, clearly after the beginning of sugar storage, but an average duration of about 45 days is widely accepted between veraison and harvest (Davies and Robinson, 1996; Sadras et al., 2008). However, studies on individual berries have revealed dramatically shorter durations, with a second growth phase of only 3 weeks on average (Bigard et al., 2019; Shahood et al., 2020; Daviet et al., 2023). This huge kinetic difference between standard samples, representative of the average population of fruits in a vineyard, and individual berries can be explained by the marked asynchrony in berry development (Rienth et al., 2016; Bigard et al., 2019; Shahood et al., 2020; Savoi et al., 2021; Daviet et al., 2023; Savoi et al., 2024; Tavernier et al., 2024). Indeed, standard samples of multiple berries collected at a given time point within a vineyard display some heterogeneity in their development, as clearly visible at the cluster level on red-skinned varieties around the phenological stage veraison. Such nonuniformity of berry development has been referred to berry asynchrony. As a consequence of this phenomenon of asynchrony, the actual rates of water and sugar accumulation in the developing berry are likely to be largely underestimated in most studies on the impact of genotype and environment on the physiology of berry ripening (Deluc et al., 2009; Leolini et al., 2019; Suter et al., 2021; Leeuwen et al., 2023). This asynchronicity bias and resulting imprecision may exceed variations caused by genotype and environment, leading to potentially erroneous conclusions.

To address pertinent metabolic fluxes inside the pericarp, there is a pressing need for non-destructive, single-berry monitoring tools that might address not only fruit expansion, such as image analysis (Daviet et al., 2023), but also the evolution of major solutes. Additionally, adapting these techniques for real-time field measurements would be crucial for studying temperature and drought tolerance, as well as genetic variability. Until now, most analytical methods used to phenotype major berry solutes have required destructive sampling and extraction of berry juice, making it impossible to monitor developmental changes in the same individual berry. Additionally, due to the considerable heterogeneity of berries within clusters, between clusters, and among plants, hundreds of fruits must be combined to produce reproducible samples. In an analogous approach to image analysis regarding the growth and coloration of individual berries (Daviet et al., 2023), non-destructive techniques, coupled with supervised models might overcome these limitations. These techniques include hyperspectral imaging and near-infrared reflectance spectroscopy (NIRS) (Ye et al., 2023).

NIRS calibration models have already been reported for the quantification of various parameters in grapevine juice or wine, such as total soluble solids (TSS), titratable acidity (TA), phenolic compounds, anthocyanins, and tannins (dos Santos Costa et al., 2019; Wyngaard et al., 2021; Rouxinol et al., 2022; Ferrara et al., 2022). These models generally reached determination coefficients (R²) above 0.85, indicating their strong prediction potential (dos Santos Costa et al., 2019; Rouxinol et al., 2022; Ferrara et al., 2022). NIRS coupled with discriminant analysis has also been used to predict the stage of grape development based on sugar content and variations in phenolic compounds during ripening (dos Santos Costa et al., 2019). Recently, NIRS calibration models have been developed to predict glucose, fructose, malic, and tartaric acids as individual compounds rather than as composite variables like TSS or TA, achieving R² values above 0.90 for sugars, 0.80 for malic acid, and 0.60 for tartaric acid (Cornehl et al., 2024). Interestingly, these last two studies were based on spectra collected on intact berries underlining the potential of NIRS for the non-destructive evaluation of their composition. Yet, both studies were conducted on detached berries under controlled laboratory conditions, and one step further toward the practical application of NIRS in the vineyard would require the development of calibration models based on spectra collected non-destructively *in situ*.

Optimal conditions for the acquisition of NIRS spectra involve clarified juices or lyophilized powders in a laboratory setting. Indeed, obtaining NIRS spectra *in situ* on plant organs like fruits presents challenges. Spectra can be distorted by fruit internal structure and granularity, as well as external factors like ambient light and temperature. To minimize these distortions, several chemometrics preprocessing methods can be applied. Standard Normal Variate (SNV) reduces light scattering and granularity effects by normalizing each spectrum to its mean vector (Barnes et al., 1989; Barnes et al., 1993). The Detrend method eliminates linear or quadratic trends in the spectra, correcting background variations and improving accuracy (Savitsky et al., 1964; Barnes et al., 1993). First and second-order derivatives enhance spectral features by eliminating background variations and superposition effects. The first-order derivative reduces baseline shifts, while the second derivative mitigates curvature variations and highlights fine structures in the spectra (Savitsky et al., 1964). By applying appropriate preprocessing methods and adapting models for real-time data collection, it is possible to enhance the accuracy and practical application of NIRS calibrations (Campos et al., 2018).

This study introduces the non-destructive *in situ* application of NIRS for monitoring the progression of major solutes during single-berry ripening. Using dedicated chemometrics techniques, we report calibration models for sugars as well as malic acid on a large set of individual berries collected across two consecutive vintages on several genotypes. We further used these models to follow sugar concentration kinetics and establish the duration of sugar accumulation in individual berries for the first time. Upon showing that ripening proceeds significantly faster than previously reported, this study provides a new proof of concept announcing a paradigm change in sampling and phenotyping strategies in grapevine, alleviating asynchronicity biases that severely limited the precision of previous studies.

## Material and Methods

### Experimental design

During summer 2021 and 2022, 10 grapevine varieties, including 9 *Vitis vinifera* L. and 1 ancient interspecific hybrid, were monitored by using a NIRS portable device (MicroNIR, OnSite-W, VIAVI) in the Pierre Galet experimental vineyard of Institut Agro Montpellier (France). This spectrometer covers a range from 950 to 1650 nm with a 6.2 nm resolution. It was equipped with the MicroNIR Tablet Probe to delimit a scanned area of 8 mm diameter on grapevine berries. A dark and a reference scan were taken every 10 minutes during the measurements. The dark scan was taken by pointing the probe through the soil direction, while the reference scan was taken using the white reference provided with the spectrometer. The NIRS monitoring was conducted around the equatorial plane of 10 berries per bunch, over 50 bunches, at 9 dates from 2021/07/09 to 2021/09/02 for Syrah, Couderc, Merlot (red varieties) and Servant (white variety), and 24 dates from 2022/07/08 to 2022/08/24 for Grenache, Mourvèdre, Morrastel, Carménère (red), Ugni blanc and Riesling (white). All the berries were randomly selected at the first date and marked with a colored marker for their monitoring at several dates.

For model calibration, some of the monitored berries were progressively sampled after spectrum acquisition at 7 dates in 2021 and 8 dates in 2022 (Table **S1**). These dates spanned berry developpement from the green stage to over-ripening. In 2021, 352 berries were collected for model development, with 307 used for training and 41 for validation. In 2022, 478 berries were collected, with 353 used for training and 125 for validation.

### Primary metabolites analysis

Sampled berries without pedicel were individually wrapped in aluminum foils and immediately frozen in liquid nitrogen. Their mass was measured in the laboratory, and they were mixed in a 50 mL falcon tube with 5x their mass of 0.25 N HCl solution. The resulting mixture was intensively shaken in the presence of a 1 cm glass bead and left to rest overnight at room temperature. In order to perform the simultaneous HPLC analysis of sugars (glucose, fructose) and organic acids (malic, tartaric, and shikimic acids), 100 µL of the juice was diluted in 1 mL of 10 mM H2SO4 solution containing 600 µM of acetic acid as an internal standard. The sample was centrifuged for 10 minutes at 13,000 rpm and then injected into the HPLC according to Rienth et al. (2016).

An expert annotation of the developmental stage of the berries was performed based on their concentration in glucose and fructose (G+F), the glucose-to-fructose ratio (G/F), and the malic-to-tartaric acids ratio (M/T). Berries were then classified as green (noted as “G”) or as ripening (noted as “R”). Details on the criteria used to annotate the berries are given in Table **S2**,

### Model development and evaluation

Data analysis and prediction models were performed using the R software (v. 4.3.2) under the RStudio (v 28.3.1) environment (R Core Team, 2023). Sugar accumulation times were calculated using Python (v. 3.10.12). The initial dataset comprised berries with both NIRS spectra and trait measurements. These traits included sugar concentration (glucose, fructose) and organic acids (malic, tartaric, and shikimic acids) measured by HPLC, as well as the mass of the berries. These data were then processed according to the workflow presented in Fig. **S1**.

### Spectra preprocessing

Different spectra preprocessings were compared after proceeding to an imputation with the maximum value to remove missing values in saturated spectra (labeled “imp”). Standard normal variate method (SNV, labeled “snv”), centered non-scaled SNV (“snv.cent”), Detrend method (“dt”) were then applied using the R package “prospectr” (v. 0.2.6, Stevens and Ramirez-Lopez, 2024). The first and second derivatives (“der1” and “der2”) were also calculated on normalized spectra (“SNV”) with the Savitzky-Golay filter implemented in the R package “signal” (v. 1.8-1, Signal developers, 2023), using smoothing window sizes of 11 and 21, respectively. Both preprocessed spectra and raw dosage data underwent separate principal component analyses (PCA).

### Initial dataset partitioning

Outliers in the initial dataset were searched using a Partial Least Squares Regression (PLSR) with a maximum of 20 components, a 5-fold cross-validation, and 50 iterations (“pls” R package v. 2.8.3, Liland et al., 2024; Seasholtz and Kowalski, 1992; Wold et al., 2001). The determination of outliers was based on the distribution of cross-validation residuals. The corresponding outliers were filtered out from the dataset before its partitioning according to the Kennard-Stone algorithm, as implemented in the “prospectr” R package v. 0.2.6 (Kennard and Stone, 1969). The partitioning enabled the selection of 1/5th of the berries, forming an external validation set, which was removed from the initial dataset to form the training set for model development. Please note that the outliers were included in the training set, and only removed from the validation set, as recommended by Metz et al. (2021).

### Developmental stage prediction

A classification model was trained using Partial Least Squares Discriminant Analysis (PLSDA) on the training set. For each pretreatment, the model underwent cross-validations (4 folds, 25 iterations). The maximal number of latent variables (LV) was set at 20, and the optimal number of LVs as well as the best spectrum pretreatment were determined as the ones minimizing the cross-validation error of classification. Model validation was performed by applying the optimal model on the previously defined external validation set. The prediction accuracy was then calculated as the proportion of correct predictions over the total number of predictions.

### Berry’s composition prediction

PLSR models were developed for the prediction of dosage data within the training set. For each spectra pretreatment, model development included an automatic detection and elimination of outliers (as previously presented), with cross-validations (5 folds, 50 repetitions). The maximum number of LVs was set at 20, and the optimal number of LVs as well as the best spectrum pretreatment were determined as the ones maximizing the cross-validation R2. The optimal model was then validated by predicting the traits on the validation set and determining the corresponding R² and RMSE.

The NIRS spectra of berries monitored throughout the entire duration of the experiment were pretreated in the same way as the spectra used for model development. For accurate enough models, traits were predicted using the optimal pretreatments and number of LVs.

### Dynamics of sugar concentration

Outliers were removed from the predicted sugar concentrations, including negative sugar concentration values and those higher than 1500 mM, obviously related to intense post-ripening shriveling (Savoi et al., 2021; Savoi et al., 2024). The resulting predictions were then plotted by the days of the year (DOY) and a 3 parameters sigmoid equation was fitted, to the evolution of sugar concentration in each berry having more than 9 valid monitoring dates, using the “NumPy” Python library v. 1.26.4 (Harris et al., 2020). These berries were monitored in 2022 on 6 kinetically extreme genotypes: Carménère, Grenache, Morrastel, Mourvèdre, Riesling, and Ugni Blanc (Suter et al., 2021). The sigmoid fits with R2 between observed and predicted values lower than 0.8 were not considered further to estimate the duration of sugar accumulation. The DOYs when the sugar concentrations reached 0.2 M (T0, the onset of berry ripening) and 1 M (T1, the point at which phloem sugar unloading typically stops in berries) were calculated, and the duration of the sugar accumulation phase time was determined by subtracting T0 from T1.

## Results

### Expert annotation of the developmental stages

A total of 830 berries were sampled immediately after spectrum acquisition, across 17 dates (9 in 2021 and 8 in 2022). An expert annotation of these berries was carried out according to their sum of glucose and fructose concentrations (G+F), and glucose-to-fructose (G/F) and malic-to-tartaric acids (M/T) ratios (Fig. **1**). More precisely, we considered 2 criteria, each corresponding to the combination of two variables: M/T and G/F, or G/F and G+F. Thresholds used for each of the criteria together with annotation rules are detailed in Table **S2**. . Among the 830 berries, 144 were annotated as “G”, 559 as “R” and 42 as “NA”. The respective evolution of “G” and “R” across time and for the two years is depicted in Fig. **1c** and **1d**, the proportion of “R” samples is larger than the proportion of “G” samples. There were also more “G” samples in 2021 than in 2022, underlining that veraison in 2022 occurred earlier than in 2021.

**Figure 1.**
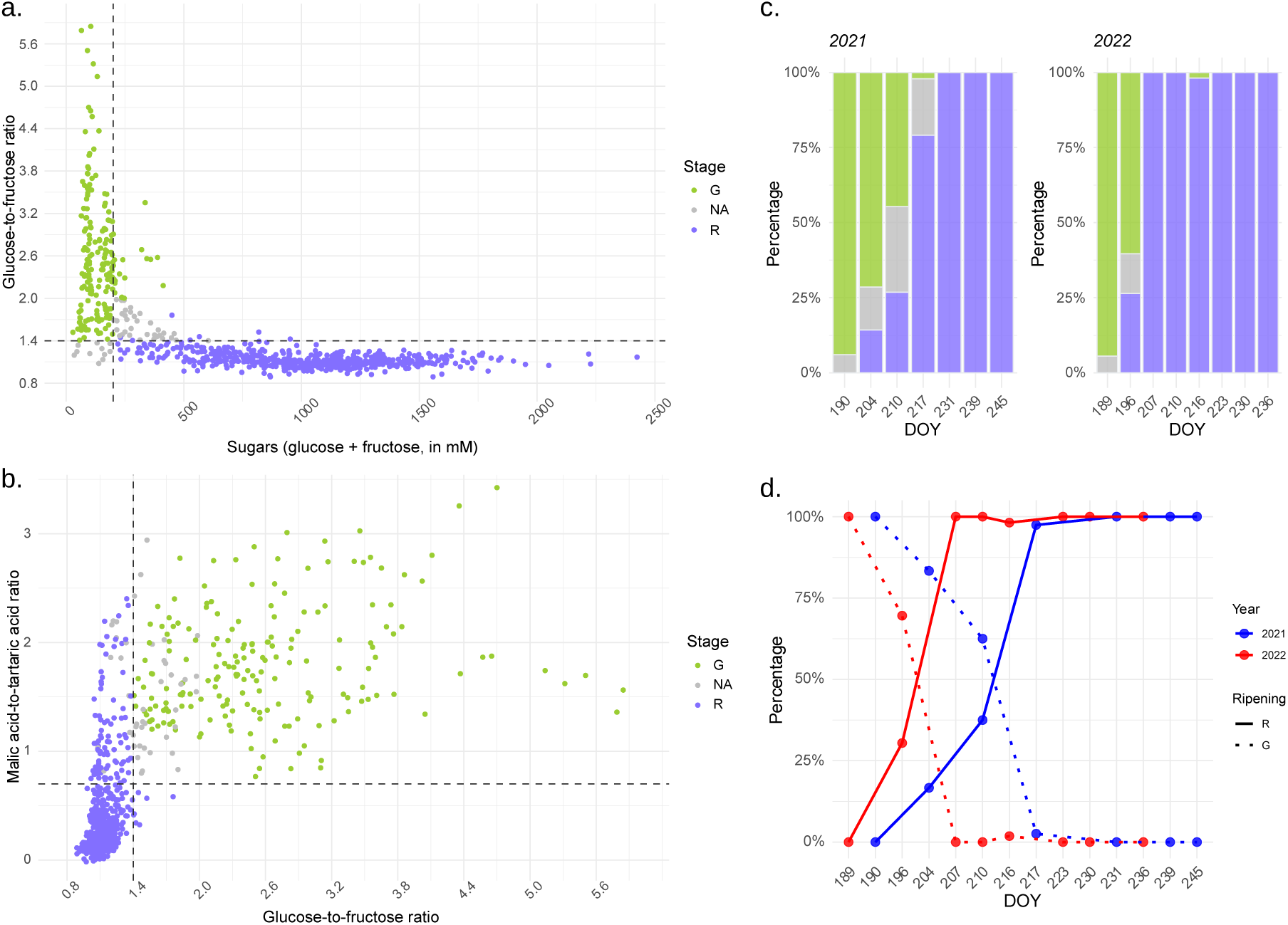
Expert annotation of the berries’ developmental stages according to their sugar concentration (glucose + fructose, in mM), glucose-to-fructose (G/F), and malic-to-tartaric acids (M/T) ratios, and distribution of the samples by their sampling date (in DOY). (**a**) The vertical dashed line was fixed at 200 mM sugars and the horizontal dashed line was at 1.4 G/F (**b**) Vertical dashed line was fixed at 1.4 G/F and the horizontal dashed line was at 0.7 M/T. (**c** and **d**) Percentage of samples annotated “before veraison” (“G”), “unidentified” and “after veraison” (“R”) by days of the year (DOY) per year.

### Exploratory analysis of reference datasets

A PCA on HPLC data highlighted a clear separation between “G” and “R” berries along PC1 (67% of explained variance), with the unidentified berries in between (Fig. **2a**). The green berries (before veraison) were, as expected, high in malic acid, G/F ratio and low in sugars, while the soft berries (after veraison) displayed high sugar concentrations (Fig. **2a** and Fig. **2b**). The second axis was more clearly associated with variations in tartaric acid concentration. The separation between “G” and “R” berries was less clear on the PCA carried out on NIR spectra, with differences depending on the pretreatments. Yet, the second derivative (der2) showed a certain separation of “G” and “R” berries (Fig. **2c**), particularly on PC2 (18.7% of explained variance).

**Figure 2.**
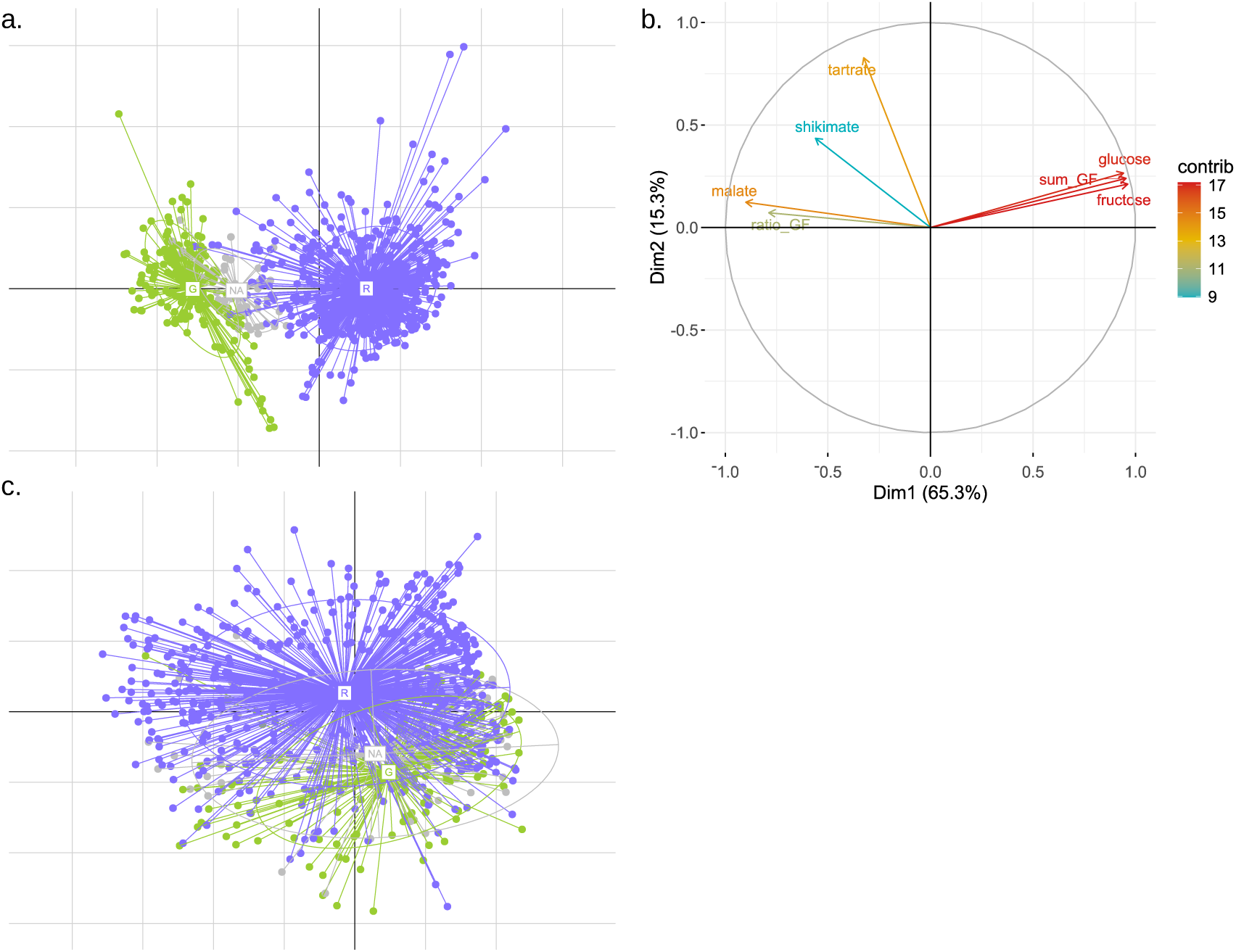
PCA on HPLC traits (a, b) and NIR spectra (c). (**a**) Projection of berries on the first PC plan with a color by the developmental stage. The x-axis represents PC1 while the y-axis represents PC2. The HPLC traits corresponded to the concentration of glucose, fructose, malic acid, tartaric acid, and shikimic acid, measured from frozen berries by HPLC, as well as the berry mass, the sum of glucose and fructose (G+F or sugars = sum_GF) and glucose-to-fructose ratio (G/F = ratio_GF). (**b**) Contribution of each HPLC trait to the first PC plan, with a color scale indicating the contribution. ( **c**) The spectra were preprocessed using the 2nd derivative (“der2”), presenting the best representation of the structure of the spectral data. As the spectra contain much more information than the more focused HPLC data, it is normal for the PCA to be noisier.

### Classification model evaluation

PLSDA was applied to predict the developmental stage of the berries. The best model within the cross-validation corresponded to the “imp” pretreatment. This model used 9 latent variables and had an average accuracy of 89.7% in the training set. It was then applied to the validation set for the prediction of expert annotated stages. The model correctly predicted all the 20 “G” annotations, reaching 100% accuracy for this class. The model also performed very well with the “R” class, correctly predicting 137 out of 139 observations (98.6% accuracy for this class). This PLSDA model is highly effective in classifying berries before or after veraison, as evidenced by a global prediction rate of 98.2%. Of the 6 “NA” observations from the validation set, the model predicted 5 as “G” and 1 as “R”.

### Berries’ composition prediction

#### Models performances in the training set

The quality of the PLSR models depended on the trait (Table **1** and Fig. **S2**). The best models were obtained for glucose and fructose, with cross-validation R² values of 0.82 and 0.76, respectively, and RMSE_cv_ values of 108 and 123 mM. For glucose, fructose, and shikimic acid, the best pretreatment was SNV, with a centered but non-scaled version for glucose. For total sugars (G+F), the best pretreatment was the second derivative (der2). As expected, the RMSE_cv_ of this model was almost twice higher (255 mM) than the one for glucose and fructose alone. The corresponding model had a R²_cv_ of 0.74. Malic acid had an R²_cv_ of 0.66 but a fairly high RMSE_cv_ value (85 mEq/L). The best model was for spectra without pretreatment, demonstrating that the raw spectra were sufficiently informative for this trait. The calibration model for berry weight also showed reasonable performance, with an R²_cv_ of 0.62 and an RMSE_cv_ of 0.3 g, and with the first derivative (der1) as the best pretreatment. However, for tartaric and shikimic acids, R²_cv_ were respectively 0.34 and 0.48, and RMSE_cv_ of 41.1 mEq/L and 479 µEq/L. While the glucose-to-fructose ratio (G/F) was predicted with reasonable accuracy, achieving an R²_cv_ of 0.7 and an RMSE_cv_ of 0.36, it seemed that a linear regression might not be the most appropriate approach for predicting this trait. Indeed, the corresponding plot indicated a curve that could be divided into two distinct linear segments (Fig. **S2**).

**Table 1.** PLSR models for berry traits. Glucose, fructose, malic acid, tartaric acid, and shikimic acid concentrations were determined by HPLC, and sugars, by the sum of glucose and fructose concentrations. Pretr.: optimal NIRS preprocessing. Nb comp: optimal number of components. R²_cv_ and R²_val_: cross-validation and validation accuracies. RMSE_cv_ and RMSE_val_: root mean square error of the model within the cross-validation and validation. RMSE is reported in the unit of measurement of the assays. Obs: number of observations used to train the model. Out: number of outliers. The validation set included 166 observations.

| Trait | Pretr. | Nb comp | $R^2_{cv}$ | $RMSE_{cv}$ | | Obs | Out | $R^2_{val}$ | $RMSE_{val}$ | |
| --- | --- | --- | --- | --- | --- | --- | --- | --- | --- | --- |
|  |  |  |  | Trait unit | g/L |  |  |  | Trait unit | g/L |
| Glucose (mM) | snv.cent | 18 | 0.82 | 108 | 19 | 630 | 30 | 0.71 | 126 | 22 |
| Fructose (mM) | snv | 19 | 0.76 | 123 | 22 | 638 | 22 | 0.64 | 132 | 24 |
| G+F (mM) | der2 | 20 | 0.74 | 255 | 87 | 640 | 20 | 0.46 | 334 | 114 |
| G/F | imp | 11 | 0.70 | 0.36 | / | 603 | 57 | 0.65 | 0.28 | / |
| Malic acid (mEq/L) | imp | 7 | 0.66 | 85 | 57 | 651 | 9 | 0.55 | 74 | 49 |
| Tartaric acid (mEq/L) | imp | 7 | 0.34 | 41 | 31 | 649 | 11 | 0.10 | 33 | 25 |
| Shikimic acid (μEq/L) | snv | 20 | 0.48 | 479 | 279 | 653 | 7 | 0.04 | 456 | 265 |
| Berry weight (g) | der1 | 14 | 0.62 | 0.3 | / | 646 | 14 | 0.43 | 0.3 | / |

#### Models v*alidation*

For the sum of sugars (G+F), the R²_val_ was 0.46, with an RMSE_val_ of 334 mM, a *c.a.* 30% increase when compared to cross-validation results (Table **1**). For malic acid, the R²_val_ was 0.55, and the RMSE_val_ was 74 mEq/L, with an RMSE decrease of about 13%. Glucose and fructose showed the smallest differences between training and validation sets. For glucose, the R²_cv_ was 0.82 compared to 0.71 for R^2^_val_, with an RMSE increase of 17%. For fructose, the R²_cv_ was 0.76 compared to 0.64 for R^2^_val_, with an RMSE increase of 9%. G/F had a R²_val_ of 0.65 and a RMSE_val_ of 0.28. Given the poor performance in the validation set of the direct prediction of the sum of sugars (G+F), an alternative approach was tested: upon adding the predicted values of glucose and fructose (sugar_cal) and comparing this value to the observed value for the sum (Fig. **3a** and **3b**). This method yielded an R²_val_ of 0.73, compared to 0.46 for the direct prediction of total sugars (G+F). Additionally, the RMSE_val_ was improved by 29% (237 mM compared to 334 mM). These results suggested that, despite their similar crude molecular formula, the aldehyde and ketone groups on, respectively C1 carbon of glucose and C2 carbon of fructose are well resolved by NIRS. This method was also applied to G/F but yielded very poor results, with a negative R²_val_ and an RMSE_val_ of 6. In fact, the current models for glucose and fructose could predict negative values, primarily in green berries that contain little or no sugar (Fig. **S2**), which may result in negative ratios, leading to poor performance.

**Figure 3.**
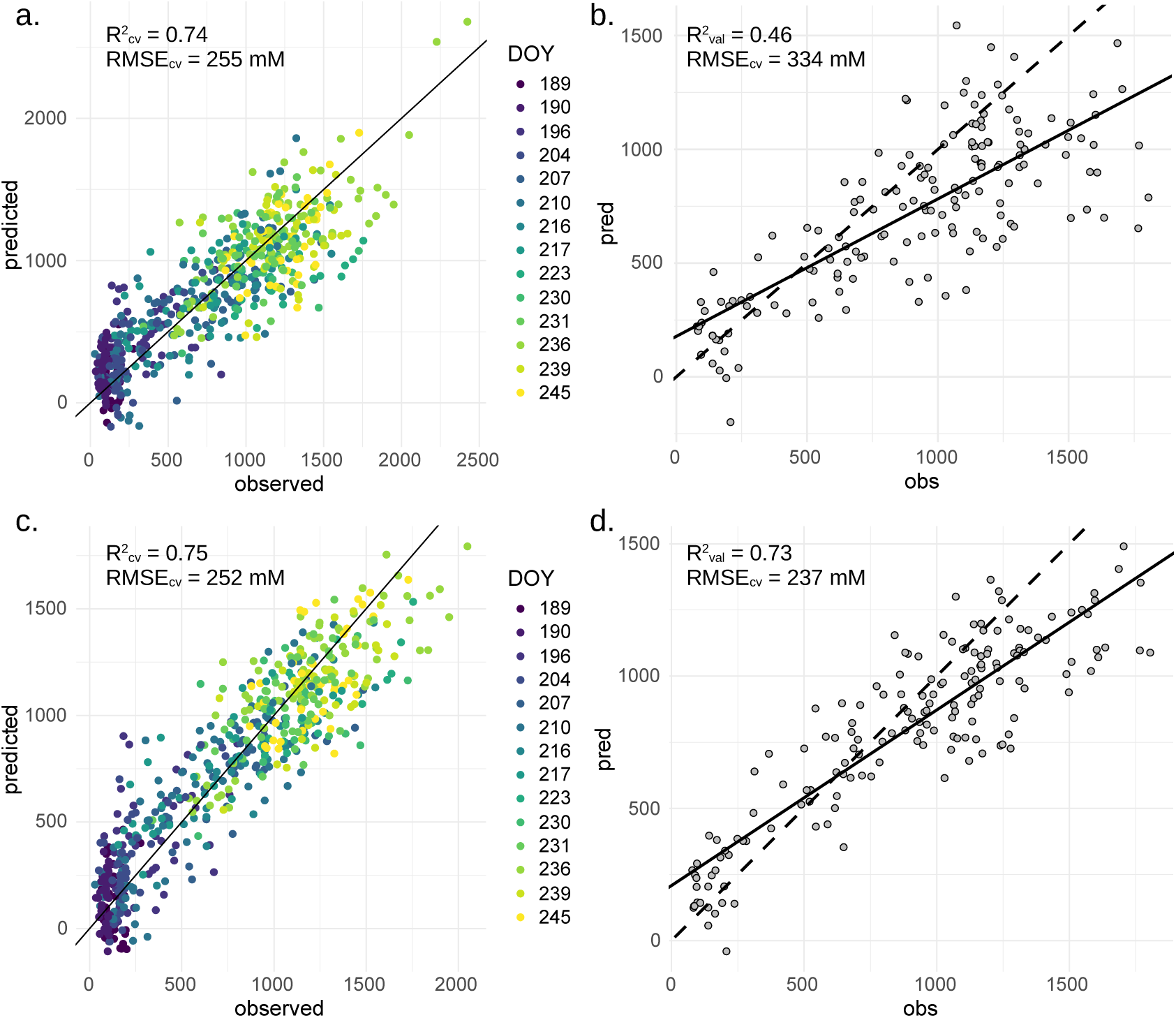
Models for predicting the concentration of sugar in individual berries from NIR spectra collected in 2021 and 2022. (**a**, **b**) Sugars represent the sum of glucose and fructose concentrations, measured by HPLC. (**a**, **c**) The results are colored by days of the year (DOY). The line represents the best-fitting line of the model. (**b**, **d**) The predictions in the validation set. The dashed line corresponds to the 1:1 line. The validation set corresponds to a separate dataset from those used to train the model (training set). (**c** and **d**) The observed Sugar_cal values correspond to the sum of the HPLC glucose and fructose assays. The predicted values correspond to the sum of the predictions for glucose and fructose models.

### Prediction of the non-sampled berries

The monitoring dataset included a total of 3,485 NIR spectra collected on 230 berries from 10 genotypes. In the absence of destructive sampling, the application of PLSR models for glucose, fructose, and malic acid can ultimately provide the specific kinetics of the sugars sum and malic acid in a given berry (Fig. **S3**). The progressive concentration/accumulation of sugars and the consumption of malic acid during the ripening phase were observable for all grape varieties. We could see, for example, that when the concentration of sugars reached its maximum from DOY 220 onwards in most grape varieties, almost all the malic acid present in the berries was consumed.

### Deciphering the evolution of sugar concentration with time in single berries

Of the 230 monitored berries, 125 had predicted data for at least 9 dates and were retained to model sugar accumulation over time. Three parameter sigmoid fits proved quite accurate with R² values ranging between 0.42 and 0.97 (median of 0.87, Fig. **4a**). Sigmoid fits with an R² higher or equal to 0.8 were considered acceptable, resulting in a total of 88 single berry kinetics (Fig. **4b**). The number of days needed to reach 1 M sugars after the onset of ripening at 0.2 M was determined from each of these fits (Fig. **4c**, Fig. **4d**). On average, it took 20 days for a berry to increase its sugar concentration from 200 to 1000 mM. The average sugar accumulation times (in days) per genotype were as follows: Morrastel = 15.7, Ugni Blanc = 19.9, Mourvèdre = 20.2, Grenache = 21.4, Carménère = 25.2, Riesling = 26.8. The durations for Carménère and Riesling were calculated with a small number of berries, 3 and 6 respectively, compared to the other genotypes (between 19 and 23 berries) (Fig. **4d**).

**Figure 4.**
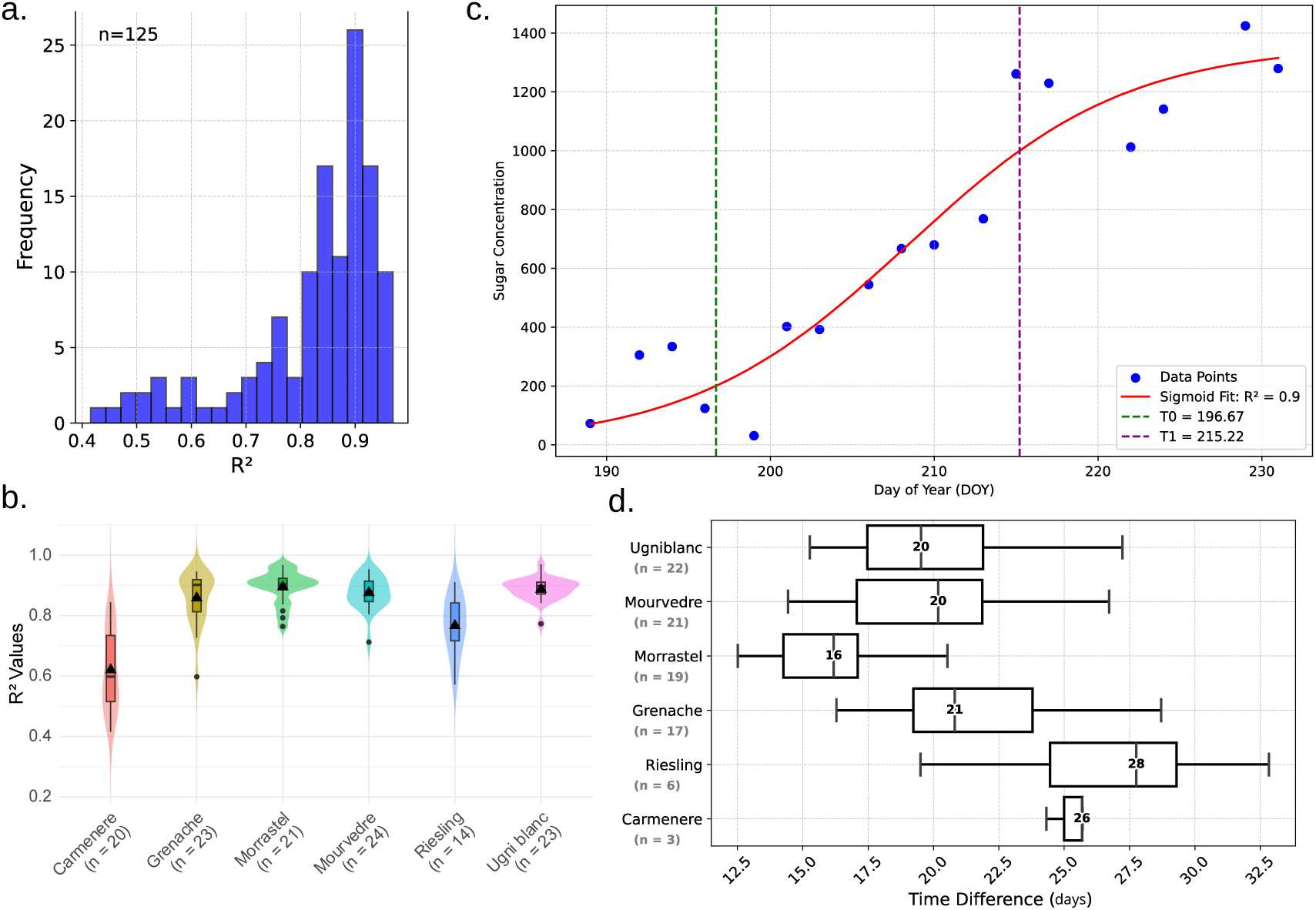
Estimation of sugar accumulation times in single berries. (a) R² distribution of sigmoid fits for sugar evolution curves for each berry monitored (125 in total). (b) R² distribution per variety. ‘n’ corresponds to the number of berries per variety. (c) An example of an estimate of the evolution of sugars in a berry of the Ugni blanc variety (dots) and its fitted sigmoid (red curve). The green dotted vertical line corresponds to T0 (= 0.2 M) and the purple line corresponds to T1 (= 1M). These two variables were used to calculate the time difference. (d) Distribution of predicted sugar accumulation times between varieties. The bold numbers inside the boxes indicate the median time difference for each genotype.

## Discussion

This study provides the first insights into the non-destructive monitoring of the evolution of the main osmotica during the ripening of individual grapes. To the best of our knowledge, although NIRS monitoring was recently applied on detached berries in laboratory conditions (Ferrara et al., 2022; Cornehl et al., 2024), no attempts were made to monitor *in situ* the progression of sugar concentration over time in single berries. Such monitoring allowed us to compare the duration of the active period of phloem unloading known as ripening, in cohorts of individual fruits from different grapevine genotypes.

In the current study, the accumulation of sugars and the consumption of malic acid during the ripening phase of individual berries were clearly observed across all grape varieties (Fig. 4 and S3). However, present models developed on field data showed lower performance compared to those based on laboratory data (Cornehl et al., 2024), as evidenced by lower R² values and higher RMSEs for glucose, fructose, malic, and tartaric acids. For example, the RMSE for glucose and fructose was 19 g/L and 22 g/L (R² = 0.82 and 0.76), compared to c.a. 9 g/L (R² = 0.92) in Cornehl et al. (2024). The RMSE for malic acid was 57 g/L (R² = 0.66) versus 3 g/L (R² = 0.84) in Cornehl et al. (2024). In line with these last authors, the impact of variety on the prediction models for glucose, fructose, and malic acid appeared negligible, alleviating the need to create a separate model for each variety (Fig. S2). Also, the fact that our model seemed to hold for both red and white varieties indicates that the NIRS prediction of the evolution of major osmotica across berry development was not deduced from their correlation with color change.

Application of NIRS to field conditions presents significant challenges since various factors can influence the spectra, including environmental conditions (Nicolaï et al., 2007; Xu et al., 2019). Sample-specific factors, such as fruit size, skin opacity, and thickness, among other internal structures influencing light penetration in the fruit, can also impact the spectra, whether taken in the field or in the laboratory (Nicolaï et al., 2007; De Oliveira et al., 2014; Campos et al., 2018; Jiang et al., 2022). For instance, the water balance of the fruit, which fluctuates throughout the day, can affect NIRS measurements. These potential issues have likely led to variations in the spectra and inaccuracies in the data, such as additive or multiplicative effects. Fortunately, some of these distortions can be corrected by preprocessing (Xu et al., 2019). Repeated handling of the fruit for spectra acquisition over time might also have damaged cuticular waxes, potentially altering both the light penetration and the physiology of the fruit. Also, NIRS measurements for monitoring fruit development typically occur during the summer, under warm conditions. To address temperature issues during measurements, Campos et al. (2018) proposed a global compensation temperature model that requires spectra acquisition at different temperatures in the training and validation sets. Further work needs to be done, to check if this method can be transferred in the field, following calibration in laboratory conditions. Finally, in previous works on detached berries, the authors carried out repeated measurements (2 to 3 scans) at different places around the berries (Ferrara et al., 2022; Cornehl et al., 2024), which might improve spectra repeatability. However, the steric hindrance of the cluster prevents the collection of multiple scans at different places around the berry in the vineyard. Nevertheless, our results showed that high-frequency measurements and curve fitting partially overcame such intrinsic noise of NIRS measurements in external conditions.

PLSR is widely used in the spectral analysis for its ability to handle high-dimensional data with complex relationships (Wold et al., 2001; Metz et al., 2021; Ryckewaert et al., 2022; Courand et al., 2022; Cornehl et al., 2024). Yet, it may not optimally predict certain traits. Deep learning techniques offer a promising avenue for improving prediction accuracy by capturing non-linear relationships within the data and extracting hierarchical representations (Vasseur et al., 2022; Houngbo et al., 2024). Fuentes et al. (2018) demonstrated the efficacy of machine learning techniques for classifying grapevine varieties using spectroscopy and morpho-colorimetry parameters, highlighting the potential of these methods to handle heterogeneous data (Danilevicz et al., 2022). Combining preprocessing and testing other dimension reduction methods could also improve model performance as tested by Ye et al. (2023). However, deep learning and neural network approaches require large amounts of data to deliver optimal results, which must be taken into account when designing experiments.

In pioneering work, Coombe & Phillips (1982) found that a single berry can reach 1.1 M sugar in two weeks only, but they considered that such an unexpectedly short duration was the consequence of a wounding artifact due to hypodermal sampling. Tedious annotation of individual softening dates and periodic destructive HPLC analysis of thousands of berries led Shahood et al. (2020) to propose a 26-day duration for sugar loading in berries of varieties Syrah and Meunier, which were synchronized according to their composition in sugars and acids. Studies on berry growth with manual measurements with calipers (Friend et al., 2009) and image analysis (Daviet et al., 2023) reported a shorter (c.a. 3 weeks) duration for the second growth phase, growth resumption being somewhat delayed with respect to softening and onset of sugar loading (Coombe and Bishop, 1980; Savoi et al., 2021). Non-destructive sugar measurement *in situ* presently shows that, as a matter of fact, the historic single berry analysis of Coombe & Phillips (1982) falls in the same duration range as our fastest ones on the variety Morrastel. The accumulation of 1 M sugars within 3 weeks in both Grenache, Ugni blanc, and Mourvèdre individual berries appears five days shorter than previously inferred in Syrah and Meunier (Shahood et al., 2020). Our sampling from the first year (2021) did not yield enough data points to properly model sugar accumulation over time. We thus carried out a second measurement campaign in 2022 with more frequent spectra acquisitions, which enabled the estimation of the sugar accumulation rate. Yet, our data also showed that these two vintages were pretty different, the veraison stage occurring earlier and faster in 2022 than in 2021. It is thus possible that our results on sugar accumulation rate might be specific to the 2022 vintage. Further dynamic NIRS data are thus needed to generalize our findings and establish the respective roles of genetic or environmental factors on the observed variation. Anyhow, the reported differences between varieties are much smaller than those reported on usual samples of unsynchronized berries representative of the global population at the vineyard scale (Table S3, Suter et al., 2021). While we do not directly prove that asynchrony is the causing factor explaining differences between our and previous works on mixture of berries, our results are in line with the hypothesis that asynchronicity bias results in an overall underestimation of solutes rate of about twofold in usual samples. Therefore, process based models of berry ripening (Dai et al., 2009; Capko et al., 2020) should be revisited and placed on a more pertinent dynamic basis. Moreover, genetic differences previously documented at population levels (Suter et al.,2021) do not translate to single berries from the same cultivars. One possible interpretation is that berry asynchronicity may change between cultivars, leading to a variable underestimation of the real rate of sugar concentration in the ripening berry according to the cultivar. Consequently, only by monitoring individual berries could real metabolic differences be identified.

## Conclusion

For the first time, using a handheld NIRS micro-spectrometer, we could monitor the evolution of grapevine berry primary metabolites, such as glucose, fructose, and malic acid, during ripening of the same berry in the vineyard. Upon alleviating severe asynchronicity biases in previous studies addressing the average population, kinetics observed on individual berries from different genotypes showed that sugar concentration proceeds twice faster than previously reported. Combining NIRS with image analysis through for instance hyperspectral imaging of the same berry will allow to calculate the net fluxes of water and major primary metabolites inside the individual fruit, as the pertinent developmental, transcriptomic, and metabolic regulatory unit. This proof of concept demonstrates these non-destructive methods deserve further optimization and automation, for identifying genetic differences and the impact of environmental conditions on the dynamics of prevailing physiological traits.

## Acknowledgments

Thanks to the Institut Agro Montpellier for the plant material and the access to the vineyard.

## Author contributions

The monitoring and collection of the grapevine berry samples were done by C. R., E. M-H, L. LC., M. T., V. S, and T. H. Sample preparation and HPLC analyses were performed by C.R. and F.T. F.T. and V.S. established and implemented the multivariate analysis workflow. C.R., F.T., and V.S. interpreted the results and drafted the manuscript. C.R. and V.S. conceived, designed the study, coordinated, and supervised the experiments. C.R. and V.S. have an equal contribution.

## Data availability

Data are available online: https://doi.org/10.57745/YGKPZA.

Scripts and code are available online: https://github.com/floratavernier/NIRS-monitoring-of-sugar-accumulation-in-single-berries.

