## Supplementary materials for "*In situ* near-infrared non-destructive monitoring of sugar accumulation reveals that single berries ripening takes only 20 days"

### Near-infrared real time non-destructive monitoring of sugar accumulation reveals that single berries ripen two times faster than previously documented on standard asynchronous samples


Flora Tavernier[1], Elias Motelica-Heino[1], Miguel Thomas[1,2], Theresa Herbold[1,2], Mengyao Shi[1], Loïc Le Cunff[1,2], Charles Romieu[1,2], Vincent Segura[1,2,*]

[1] UMR AGAP Institut, Univ Montpellier, CIRAD, INRAE, Institut Agro, F-34398 Montpellier, France.
[2] *Geno-Vigne, IFV-INRAE-Institut Agro, F-34398, Montpellier, France.*


**Table S1. Dataset description.**

| Variety | Year | Skin color | HPLC and NIRS (model) | | NIRS monitoring | |
|---|---|---|---|---|---|---|
| | | | Number of samples | Number of dates | Number of spectra | Number of dates |
| Couderc | 2021 | Red | 87 | 7 | 189 | 9 |
| Merlot | 2021 | Red | 98 | 7 | 252 | 9 |
| Servant | 2021 | White | 83 | 7 | 144 | 9 |
| Syrah | 2021 | Red | 84 | 7 | 216 | 9 |
| Carménère | 2022 | Red | 86 | 8 | 452 | 20 |
| Grenache | 2022 | Red | 80 | 8 | 487 | 20 |
| Morrastel | 2022 | Red | 68 | 8 | 442 | 19 |
| Mourvèdre | 2022 | Red | 84 | 8 | 431 | 19 |
| Riesling | 2022 | White | 77 | 8 | 401 | 20 |
| Ugni blanc | 2022 | White | 83 | 8 | 471 | 19 |
| **Total of samples and NIRS monitoring** | | | **830** | **15** | **3,485** | **25** |

**Table S2. Rules for berry annotations.** Two criteria, each including the combination of 2 traits, were used to annotate the berries. A berry was considered to be before veraison when it was annotated as « G » for both criteria or « G » for one criterion and « NA » for the other. Conversely, a berry was considered te be after veraison when it was annotated « R » for both criteria or « R » for one criterion and « NA » for the other. In any other case (« G » and « R » or « NA » and « NA »), the berry was annotated as NA.

| | Criterion 1 | | Criterion 2 | |
|---|---|---|---|---|
| | M/T | G/F | G/F | G+F |
| Before Veraison (G) | > 1.4 | >0.7 | >2 | <200 |
| After Veraison (R) | <1.4 | <0,7 | <2 | >200 |
| NA | > 1.4 | <0.7 | >2 | >200 |
| | <1.4 | >0.7 | <2 | >200 |

**Table S3. Comparison of sugar accumulation time on Single Berries (this work) and usual samples (Suter et al., 2021) on the same range of cultivars.**

| | Single berry median (days) | Suter *et al.*, 2021 (days) | Suter *et al.*, 2021 / Single berry median |
|---|---|---|---|
| Ugni blanc | 20 | 52 | 2.6 |
| Mourvedre | 20 | 57 | 2.9 |
| Morrastel | 16 | 41 | 2.6 |
| Grenache | 21 | 34 | 1.6 |
| Riesling | 28 | 32 | 1.1 |
| Carmenere | 26 | 41 | 1.6 |
| **Mean** | **21.8** | **42.8** | **2.0** |

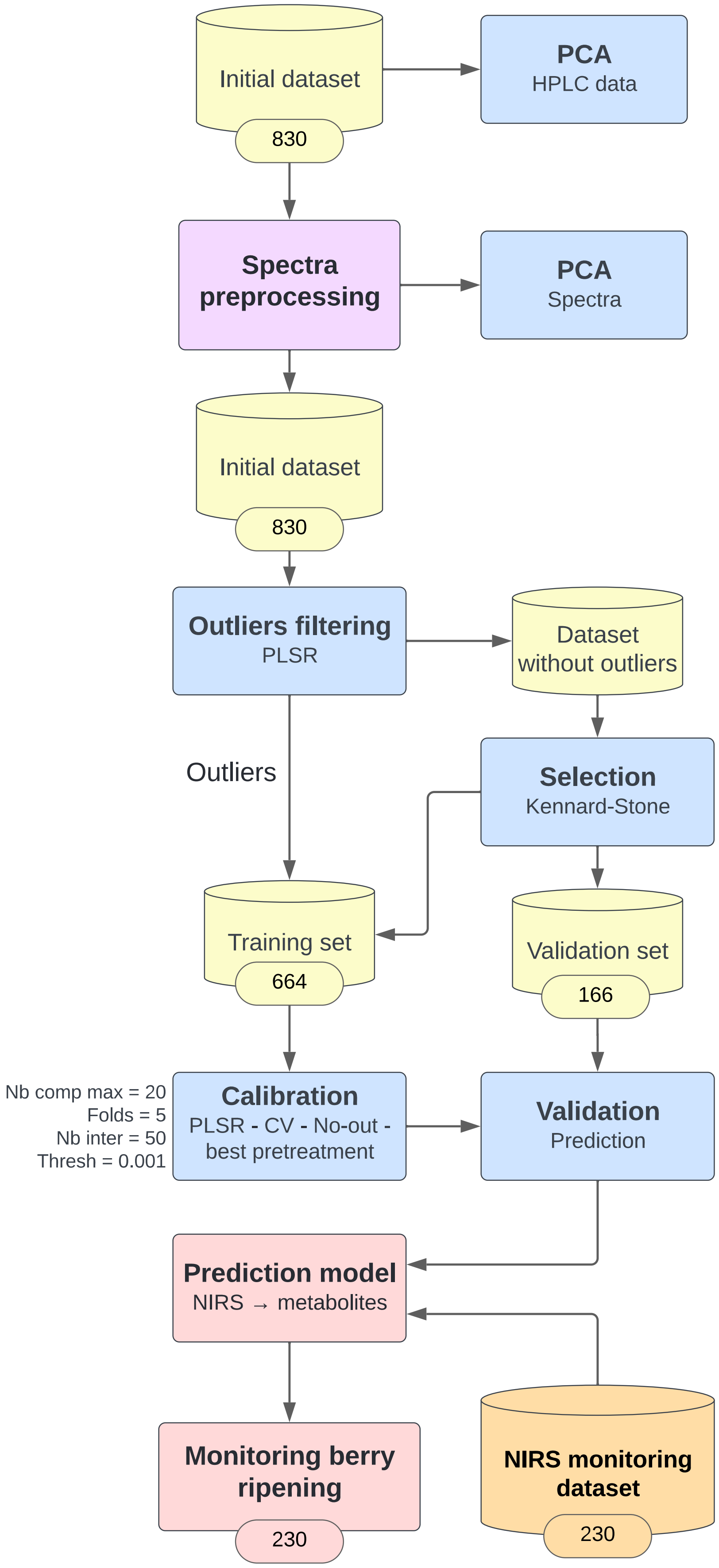


**Figure S1. Data processing and prediction model workflow.** The initial dataset included NIRS and trait measurements from 830 grapes, covering sugars (glucose, fructose), organic acids (malic, tartaric, shikimic acids) measured by HPLC, and berry mass. Spectra were preprocessed using chemometrics methods before being analyzed using PCA. Outliers were filtered out using PLSR and the validation set (n = 166 berries) was determined on the resulting data using the Kennard-Stone algorithm. This validation set was then removed from the initial dataset to create the training set (n = 664 berries). PLSR models were trained with outlier detection and cross-validation (maximum of 20 latent variables, 5-fold, 50 repetitions). The best spectral pretreatment and the optimal number of latent variables were determined for each trait with the cross-validation $R^2$. The resulting model was then applied to the validation set. Finally, the validated models were applied to the NIRS monitoring dataset (without HPLC measurements) to determine berry composition

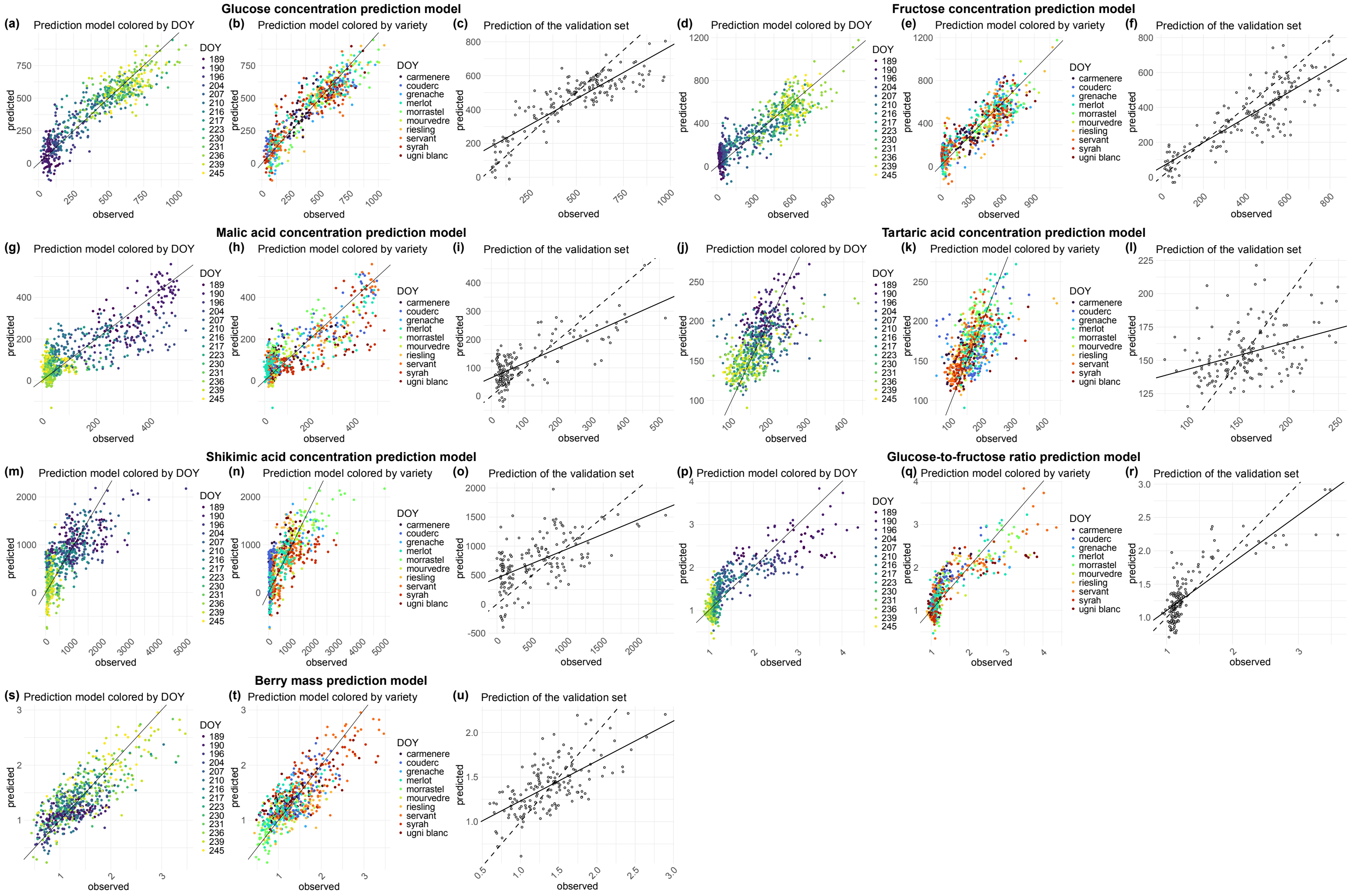


**Figure S2. Prediction models of the glucose (mM), fructose (mM), malic acid (mEq/L), tartaric acid (mEq/L), shikimic acid (µEq/L) and berry mass (g) from NIR spectra in 2021 and 2022.** The model carries out a series of cross-validations (cv) between the measured data (in x) and the predicted data (in y) before generating the final graphs, accompanied by the $R^2$, which corresponds to the accuracy of the model, and the RMSE, which corresponds to the order of error of the model in the unit of measurement of the assays (mM for sugars and mEq/L for malic acid). The results of the cross-validation (cv) are coloured by days of the year (DOY) (**a**, **d**, **g**, **j**, **m**, **p**) and by variety (**c**, **f**, **i**, **l**, **p**, **r**). (**b**, **e**, **h**, **k**, **n**, **q**) Predictions of the validation set. The validation set corresponds to a separated dataset from those used to train the model (training set).

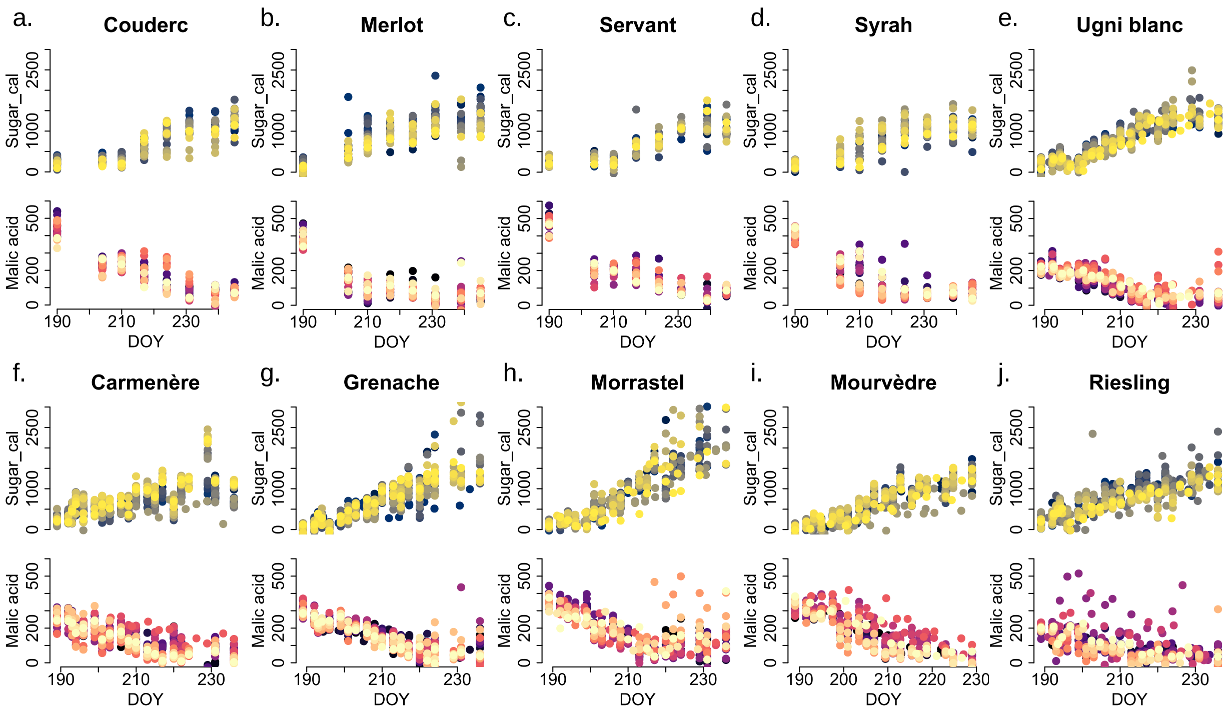


**Figure S3. Predictions of changes in sugars (sugar_cal, in mM) and malic acid concentration (in mEq/L) during ripening of single berries.** Each color represents the evolution of the same individual berry over time, monitored non-destructively by NIRS using models calibrated from HPLC data. 2021 grape varieties: Servant, Couderc, Merlot and Syrah. Grape varieties in 2022: Carmenère, Grenache, Ugni Blanc, Morrastel, Risling and Mourvèdre. Sugar_cal corresponds to the sum of glucose and fructose predicted by separate models.
